# Prioritized memory access explains planning and hippocampal replay

**DOI:** 10.1101/225664

**Authors:** Marcelo G. Mattar, Nathaniel D. Daw

## Abstract

To make decisions, animals must evaluate outcomes of candidate choices by accessing memories of relevant experiences. Yet little is known about which experiences are considered or ignored during deliberation, which ultimately governs choice. Here, we propose a normative theory to predict which memories should be accessed at each moment to optimize future decisions. Using nonlocal “replay” of spatial locations in hippocampus as a window into memory access, we simulate a spatial navigation task where an agent accesses memories of locations sequentially, ordered by utility: how much extra reward would be earned due to the computation enabling better choices. This prioritization balances two desiderata: the need to evaluate imminent choices, vs. the gain from propagating newly encountered information to predecessor states. We show that this theory offers a unifying account of a range of hitherto disconnected findings in the place cell literature such as the balance of forward and reverse replay, biases in the replayed content, and effects of experience. Accordingly, various types of nonlocal events during behavior and rest are re-interpreted as instances of a single choice evaluation operation, unifying seemingly disparate proposed functions of replay including planning, learning and consolidation, and whose dysfunction may underlie pathologies like rumination and craving.

## 1 Introduction

A hallmark of adaptive behavior is the effective use of experience to maximize reward^1^. In sequential decision tasks such as spatial navigation, actions can be separated from their consequences in both space and time. Anticipating these consequences, so as to choose the best actions, thus often requires integrating multiple intermediate experiences from pieces potentially never experienced together^2,3^. For instance, planning may involve sequentially retrieving experiences to prospectively compose a series of possible future situations^4–6^. Recent theories suggest that humans and animals selectively engage in such prospective planning as appropriate to the circumstances, and that omitting such computations can lead to phenomena of habits and compulsion^7–12^. However, by focusing only on whether or not to deliberate about the immediate future, these theories largely fail to address which of the many possible experiences to consider in this evaluation process, which ultimately governs which decisions are made.

In addition to prospective planning, behavioral and neuroimaging data suggest that actions can also be evaluated by integrating experiences long before decisions are needed. In humans, for instance, decisions in sequential tasks can indeed be predicted from prospective reinstatement of future paths at choice time^5^. However, this effect is hardly unique: Future decisions can also be predicted from analogous neural reinstatement when relevant information is first learned^13^ and also during subsequent rest^14,15^ (Fig. 1a). This is consistent with the intuition that we are generally able to think about the past, the present, or the future. Yet, it further highlights the selection problem: If actions can be evaluated long before they are needed, which experiences should the brain consider at each moment, and in what order, to set the stage for the most rewarding future decisions? These questions are likely central not only to healthy decisions, but to a variety of dysfunctions including hallucations, craving, and rumination^16,17^. Addressing them requires a new, more granular theory predicting the specific patterns in which individual memories should be accessed during deliberation. Such patterns should include prospective planning as a special case, but should generalize to additional ways of computing values by accessing memories in different orders, at different times, as observed experimentally.

**Figure 1:**
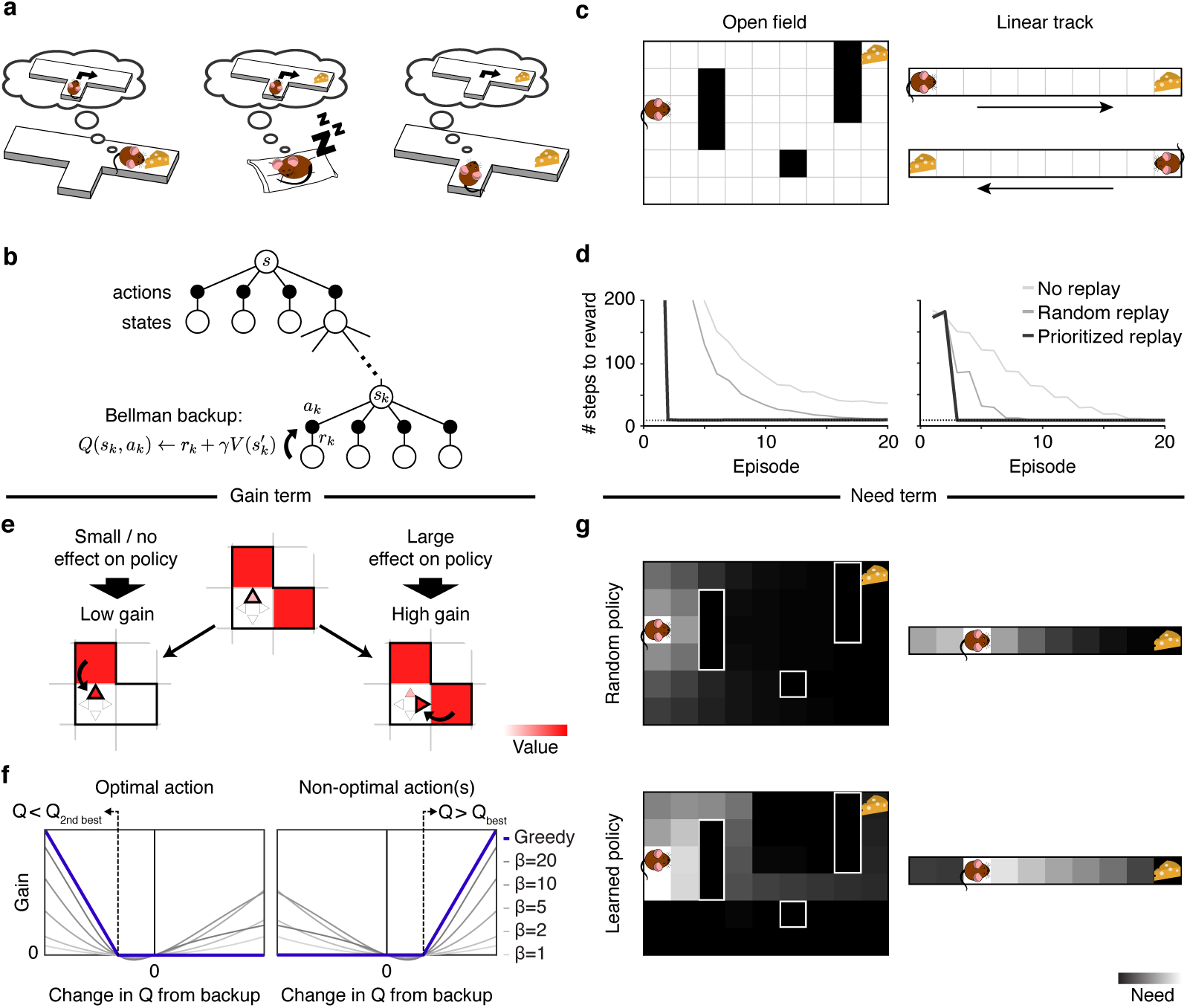
A rational model of prioritized memory access. *(a)* Three ways an agent might learn, through sequential memory access, the relationship between actions and rewards: *Left*: when reward is first encountered, through reverse reactivation; *Center*: during sleep or rest through “offline” reactivation of the sequence; *Right*: prior to choice, by prospective (forward) activation. The latter case is the most commonly envisioned in theories of model-based deliberation, but replay of all three sorts exist, and human neuroimaging evidence suggests that all can support decisions. *(b)* A schematic of a Bellman backup where the reactivation of a nonlocal experience *e_k_* = (*s_k_*, *a_k_*, *r_k_*, *s*′*_k_*) propagates the one-step reward *r_k_* and the discounted value of *s′_k_* to the state-action pair (*s_k_,a_k_*). *(c)* Grid-world environments simulated. *Left*: A two-dimensional maze (Sutton’s DYNA maze^34^) with obstacles. *Right*: a linear track simulated as two disjoint segments (to reflect the unidirectionality of the hippocampal place code in linear tracks) with rewards in opposite ends. *(d)* Performance of a greedy agent in the two simulated environments. Replaying experiences according to the proposed prioritization scheme speeds learning compared to learning without replay or with replay of randomly ordered experiences. *Left*: Open field; *Right*: linear track. Dotted lines represent optimal performance. *(e)* The gain term for updating the value of a target action in a target state quantifies the expected increase in reward following a visit to the target state. *Left*: if the best action is updated with a higher value, the policy changes little/nothing, resulting in a small/zero gain. *Right*: if a non-optimal action is updated with value higher than the best action’s value, the policy in the corresponding state changes, resulting in a large gain. Here, squares represent states, triangles represent actions, and the arrow represents a Bellman backup which updates the value of an action. The highlighted triangle represents the action with highest estimated value. *(f)* For a greedy agent (one who always chooses the best action; blue line), the gain is positive either when the best action is found to be worse than the second best action (*left*, changing the policy to disfavor it) or when a suboptimal action is found to be the best action (*right*, changing the policy to favor it). In both cases, the gain increases depending how much better the new policy is. Otherwise, the gain is zero, reflecting no effect in the policy. For a non-greedy agent (one who sometimes chooses random exploratory actions; thin gray lines), changes in *Q*-values that do not change the best action can nonetheless affect the degree of exploration, leading to nonzero gain (*β*: softmax inverse temperature parameter). Notice that a perfectly symmetric gain around zero amounts to prediction error. *(g)* The need term for a particular target state corresponds to its expected future occupancy, measuring how imminently and how often reward gains will be harvested there. This is shown as a heat map over states, and also depends on the agent’s future action choice policy, e.g. *Top*: Random policy (initially). *Bottom*: Learned policy (following training).

Empirically, a valuable window into patterns of memory access is offered by the hippocampus, a structure known to play a critical role in memory for events and places^18,19^. During spatial navigation, neural activity recorded from hippocampal place cells typically represents the animal’s spatial position, though it can also represent locations ahead of the animal during movement pauses^20–22^. For instance, during “sharp wave ripple” events, activity might progress sequentially from the animal’s current location towards a goal location^21,22^. These “forward replay” sequences predict subsequent behavior and have been suggested to support a planning mechanism that links actions to their deferred consequences along a spatial trajectory^22^. However, this pattern is also not unique: Remote activity in the hippocampus has long been known also to represent locations behind the animal^21,23–26^, and even altogether disjoint, remote locations (especially during rest or sleep^27,28^; Fig. 1a). Collectively, these three patterns (forward, reverse, and offline replay) parallel the circumstances, discussed above, in which reinstatement in humans was later shown to predict choice. The various different patterns of hippocampal replay and their occurrence in different circumstances have been suggested to support a range of distinct functions such as planning^20,22^, learning through credit assignment^23,26,29^, memory retrieval^30,31^, consolidation^30,32^, and forming and maintaining a cognitive map^25,33^. Yet, we still lack a theory describing how these various functions of replay come together to promote adaptive behavior, and predicting which memories are replayed at each time and in which order.

To address this gap, we develop a normative theory to predict not just whether but which memories should be accessed at each time to enable the most rewarding future decisions. Our framework, based on the DYNA reinforcement learning (RL) architecture^34^, views planning as learning about values from remembered experiences, generalizing and re-conceptualizing work on trade-offs between model-based and model-free controllers^7,8^. We derive, from first principles, the utility of retrieving each individual experience at each moment to predict which memories a rational agent ought to access to lay the groundwork for the most rewarding future decisions. This utility is formalized as the increase in future reward resulting from such memory access and is shown to be the product of two terms: a gain term which prioritizes states behind the agent when an unexpected outcome is encountered; and a need term which prioritizes states ahead of the agent that are immediately relevant. Importantly, this theory at present investigates which experience among all would be most favorable in principle; it is not intended as (but may help point the way toward) a mechanistic or process-level account of how the agent might efficiently find them.

To test the implications of our theory, we simulate a spatial navigation task in which an agent generates and stores experiences which can be later retrieved. We show that an agent that accesses memories sequentially and in order of utility produces patterns of sequential state consideration that resemble place cell replay, reproducing qualitatively and with no parameter fitting a wealth of empirical findings in this literature including (i) the existence and balance between forward and reverse replay; (ii) the content of replay; and (iii) effects of experience. Thus, we propose the unifying view that all patterns of replay during behavior, rest, and sleep reflect different instances of a more general state retrieval operation that integrates experiences across space and time to propagate value and guide decisions. This framework formalizes and unifies aspects of the various putatively distinct functions of replay previously proposed, and may shed light onto related psychiatric disorders including craving, hallucinations, and rumination.

## 2 Results

We consider a class of sequential decision tasks where an agent must decide in each situation (state; e.g. a location in a spatial task) which action to perform with the goal of maximizing its expected future reward. The optimal course of action (policy) consists of selecting the actions with highest expected value. The value of an action (*Q*-value) is defined as the expected discounted future reward from taking that action and following the optimal policy thereafter. Optimal decision making, therefore, requires the agent to estimate action values as accurately as possible for maximizing total reward.

We address how best to order individual steps of computation, known as *Bellman backups* (Fig. 1b), for estimating an action’s value. A single Bellman backup updates the estimate of the future value of taking a particular “target” action in some state, by summing the immediate payoff received for the action with the estimated future value of the successor state that follows it. This backup operation is fundamental for predicting future reward in RL, because it propagates information about reward to states and actions that precede it. Bellman backups can be applied action-by-action during ongoing behavior to allow the agent to learn from experienced states and rewards; this corresponds the standard update rule for “model-free” temporal difference (TD) learning, as thought to be implemented in the brain by dopaminergic prediction errors^35^. Our account includes this sort of learning from experienced events as a special case, but also allows for additional Bellman backups to be performed to update estimates for target states and actions that are not currently being experienced (Fig. 1a,b). In these cases, the resulting reward and successor state are given by remembered or simulated experiences, but the learning rule is otherwise the same. In computer science, this approach is known as the DYNA framework^34^. We refer to the information processed in a nonlocal backup as a “memory” — a target state and action, and the resulting reward and successor state. However, the same approach applies regardless of whether this information is a retrieved record of an individual event (like an episodic memory), or instead a simulated experience (a sample drawn from a learned “world model” of the overall statistics of state transitions and rewards, more like a semantic memory). These two representations are largely the same in the present work because we simulate only fixed, deterministic tasks (Fig. 1c). Importantly, because this process can compose behavioral sequences of simulated experience from pieces not experienced together, it can discover consequences missed by TD learning, which evaluate actions only in terms of their directly experienced outcomes (Fig. 1d)^35^.

Stringing together multiple backup operations over a sequence of states and actions computes expected value over a trajectory. Thus, the value of an action — the expected cumulative discounted reward that will follow its execution — can be sampled by adding up expected immediate rewards over a trajectory of one or more forward steps, plus any additional value expected forward from the last state considered. This is known as an *n*-step Bellman backup or a rollout, and can be composed from a series of one-step backups using a learning mechanism called eligibility traces^1^. Similarly, value information can be propagated backwards along a trajectory (i.e. from a destination state to each of a series of predecessors) by chaining successive one-step backups in the reverse direction. Both of these patterns (forward and reverse value propagation) have precedent in different computer science methods (e.g. Monte Carlo tree search and Prioritized Sweeping). Indeed, various existing “model-based” algorithms for computing values from a world model amount to a batch of many such backup operations, performed in different orders^1,2^. A major goal of our theory is to provide a principled account of when each pattern is most useful.

To analyze the optimal scheduling of individual steps of value computation, we derive the instantaneous utility of every possible individual Bellman backup: the expected increase in future reward that will result if the backup is executed (see Methods for formal derivation). The intuition is that an individual backup, by changing action values, has the potential to improve the choice policy at a target state, leading to better rewards if that state is ever visited. Thus, the utility of a backup can be intuitively understood as the increase in reward at the target state multiplied by the expected number of times the target state will be visited — the product of a *gain* and a *need* term, respectively. The gain term quantifies the net increase in discounted future reward expected from a policy change at the target state — that is, it measures how much more reward the agent can expect to harvest following any visit to the target state, due to what it learns from the update (Fig. 1e). Notice that this value depends on whether the update changes the agent’s policy, meaning that (in contrast to other prioritization heuristics considered in AI;^36–38^), the theory predicts asymmetric effects of positive and negative prediction errors due to their differential effect on behavior (Fig. 1f). To determine priority, the gain term is multiplied by the need term, which quantifies the number of times the agent is expected to harvest the gain by visiting the target state in the future. Here, earlier visits are weighted more heavily than later visits due to temporal discounting. This weighting implies that the need term prioritizes the agent’s current state, and others likely to be visited soon (Fig. 1g). The utility of a backup, therefore, depends simultaneously on gain and need. Thus, a backup that has no effect on behavior has zero utility even if the target state is expected to be visited in the future (because it has zero gain, despite high need). Similarly, the utility of a backup is zero if a state is never expected to be visited again, even if this backup would greatly impact behavior at the that state (because it has zero need, despite high gain). Notice that utility is computed separately for each individual backup. This “myopic” view neglects the possibility that a backup may harvest additional gains by setting the stage for other, later backups.

To test the implications and properties of this theory, we simulate an optimal agent’s behavior in a spatial navigation task (a “grid-world”) where states are locations in the environment and an action is a step of movement in one of the four cardinal directions. We simulate two distinct spatial environments (Fig. 1c). First, we simulate a linear track where the agent shuttles back and forth to collect rewards at the ends, a task widely used in studies of hippocampal replay (Fig. 1c, *right*). Second, we simulate a two-dimensional field with obstacles (walls) where the agent needs to move toward a reward placed at a fixed location, a task used extensively in previous RL studies using the DYNA framework^1,37^ (Fig. 1c, *left*). In both cases, the agent learns which actions lead to a reward by propagating value information through Bellman backups. To promote continuous learning, we add a small amount of noise to each encountered reward.

We assume that when the agent is paused (here, before starting a run and upon receiving a reward), it may access nonlocal memories, and that it does so sequentially and in order of utility. By repeating this reactivation sequentially, value information can be propagated along spatial trajectories that may have never been traversed continuously by the agent. In particular, value information can be propagated backwards by chaining successive backups in the reverse direction, or forwards by chaining successive backups in the forward direction. The latter case is achieved by allowing the agent to look one step deeper into the value of an action — i.e., we consider the utility of all individual backups and, in particular, one that extends the previous backup with one extra state and updates the values of all actions along a trajectory. This approach, which follows from first principles, results in symmetric forward and reverse updates that have comparable effects along all the states of a trajectory.

We also assume that this local operation is accomplished in the brain by place cell activity at the target location, allowing predictions of patterns of replay. Theories of model-free reward learning in navigational tasks typically assume that a hippocampal location representation is the input to a learned value mapping in striatum, updated by dopaminergic prediction error^39^. Trajectory replay in the hippocampus, which drives activation and plasticity throughout this system, is thus a potential substrate for value learning^40–42^ — effectively, by driving the dopaminergic value learning system using simulated rather than actual experiences. However, the brain may resort to different biological mechanisms to propagate value along replayed trajectories. The present work aims only to expose the principles governing how such mechanisms should behave.

### 2.1 Memory access and learning

Our first prediction was that prioritized memory access accelerates learning in a spatial navigation task. In both environments, we contrasted an agent that accesses memories in a prioritized order with a baseline model-free agent that learns only by direct experience, and with an agent that simulates experiences drawn at random (original DYNA^34^). In all cases, the number of steps required to complete an episode (the time between receiving two rewards, equivalent to a trial in a typical experiment) is gradually reduced as the agent learns the task. Learning with prioritized experience replay, however, progresses faster due to the agent’s ability to rapidly propagate value information along relevant trajectories (Fig. 1d). Notice that our theory predicts that a model-free agent is nonetheless able to learn this type of task, albeit in a slower fashion, in line with empirical findings where the disruption of replay slows down learning without abolishing it completely^31^.

### 2.2 Context-dependent balance between forward and reverse sequences

A major prediction of our theory is that patterns of memory access are not random, but often involve adjacent locations to propagate value across spatial locations iteratively. In our simulations, we observed that, as in hippocampal recordings, replayed target states typically followed continuous sequences in either forward or reverse order (Fig. 2). In the model, this is because backup operations tend to produce situations that favor adjacent backups. In particular, our theory predicts two predominant patterns of backup, driven by the two terms of the prioritization equation.

**Figure 2:**
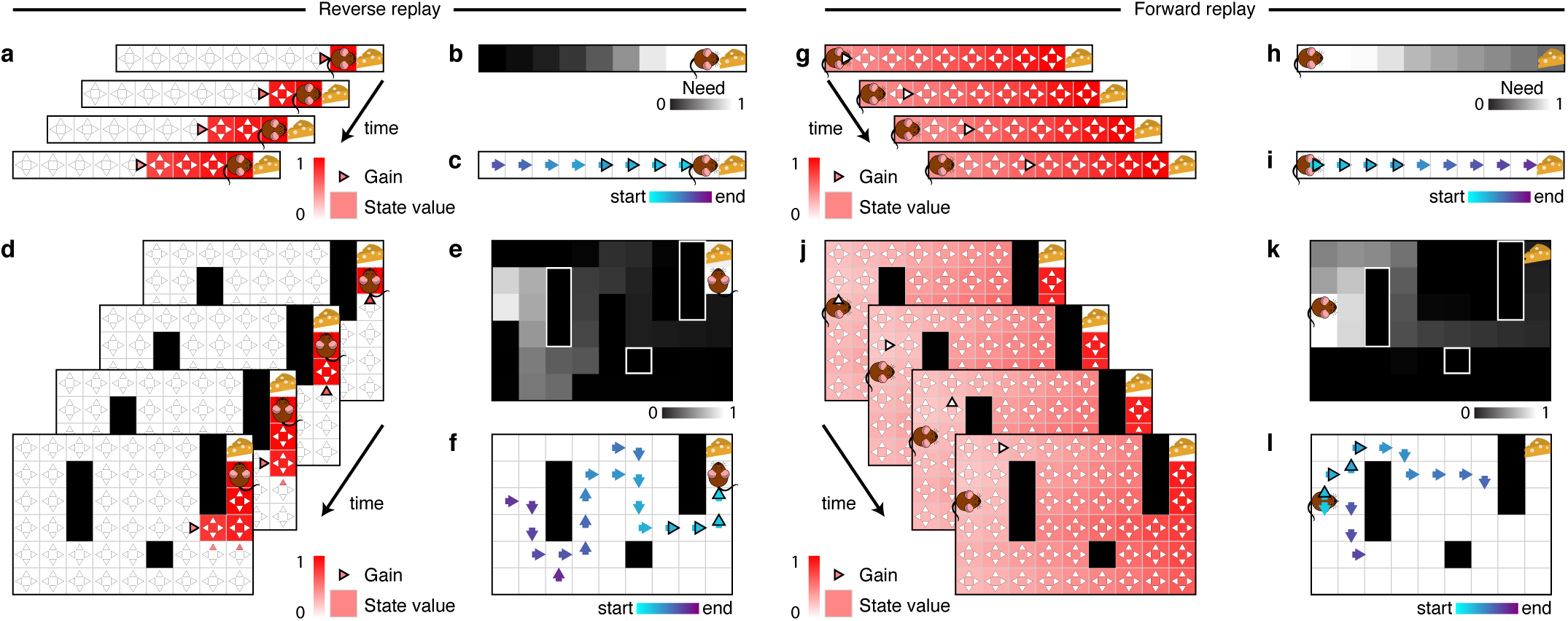
Replay produce extended trajectories in forward and reverse directions. *(a-f)* Example of reverse replay. *(g-l)* Example of forward replay. *(a,d)* Gain term and state values. Notice that the gain term is specific for each action (triangles), and that it may change after each backup due to its dependence on the current state values. Replay of the last action executed before finding an unexpected reward often has a positive gain because the corresponding backup will cause the agent to more likely repeat that action in the future. Once this backup is executed, the value of the preceding state is updated and replaying actions leading to this updated state will have a positive gain. Repeated iterations of this procedure leads to a pattern of replay that extends in the reverse direction. The highlighted triangle indicates the action selected for value updating. *(g,j)* If gain differences are smaller than need differences, the need term dominates and sequences will tend to extend in the forward direction. *(b,e,h,k)* Need term. Notice that the need term is specific for each state and does not change after each backup due to being fully determined by the current state of the agent. The need term prioritizes backups near the agent and extends forwards through states the agent is expected to visit in the future. In the field, the need term is also responsible for sequences expanding in a depth-first manner as opposed to breadth-first. *(c,f)* Example reverse sequences obtained in the linear track (c) and open field (f). *(i,l)* Example forward sequences obtained in the linear track (i) and open field (l). Notice that forward sequences tend to follow agent’s previous behavior but may also find new paths towards the goal.

First, when an agent encounters a prediction error, this produces a large gain term behind the agent (Fig. 2a-f), reflecting the gain from propagating the new information to potential predecessor states. Following such a backup, gain now also favors, recursively, propagating the information toward that state’s predecessors, and so on. In this case, sequences tend to start at the agent’s location and move backwards towards the start state (Fig. 2c,f). Because the need term is largest for states the agent expects to visit next (Fig. 2e), and since it includes a transition from the reward to the predicted start state, prioritized backups often extend backwards in a depth-first manner even in a 2D environment (Fig. 2f). The depth-first pattern is a reflection of the agent’s expectation that it will return to the reward in the future following a trajectory similar to that followed in the past, in contrast to a breadth-first pattern observed in alternative prioritization heuristics that do not include a need term.^36–38^.

The need term, in turn, tends to be largest in front of the agent (Fig. 2g-l); when it dominates, sequences tend to start at the agent’s location and move forward toward the reward location (Fig. 2i,l). They tend to iterate forward because following a forward sequence of *n* steps, an additional step can extend it to an *n* +1-step backup that carries information about each preceding action. This pattern is observed whenever the utility of looking one step deeper into the value of the actions along the route is sufficiently high.

These findings largely reproduce the different patterns of reactivation observed in both humans and rodents, suggesting that they can be explained via the same prioritized operation applied in different situations.

The model thus predicts when different patterns of backup (driven by gain and need) are likely to occur. To quantify these observations in simulation, we classified each individual backup as *forward* or *reverse* by examining the next backup in the sequence. When a backed-up action was followed by a backup in that action’s resulting state, it was classified as a *forward*. In contrast, when the state of a backup corresponded to the outcome of the following backed-up action, it was classified as *reverse*. Backups that did not follow either pattern were not classified in either category. We then followed conventional empirical methods for identifying hippocampal replay events and assessed the significance of all consecutive segments of forward or reverse backups of length five or greater with a permutation test^21,23,24^.

In line with rodent electrophysiology data on the linear track, we observed that replay (driven by the need term) extended in the forward direction before a run (Fig. 3a, *left*), providing information relevant for evaluating future trajectories. In contrast, replay extended in the reverse direction upon completing a run (driven by the gain term, Fig. 3a, *right*), providing a link between behavioral trajectories and their outcomes. Very few reverse sequences were observed prior to the onset of a run, and very few forward sequences were observed upon completion of a run, in line with previous findings^21^ (Fig. 3b).

**Figure 3:**
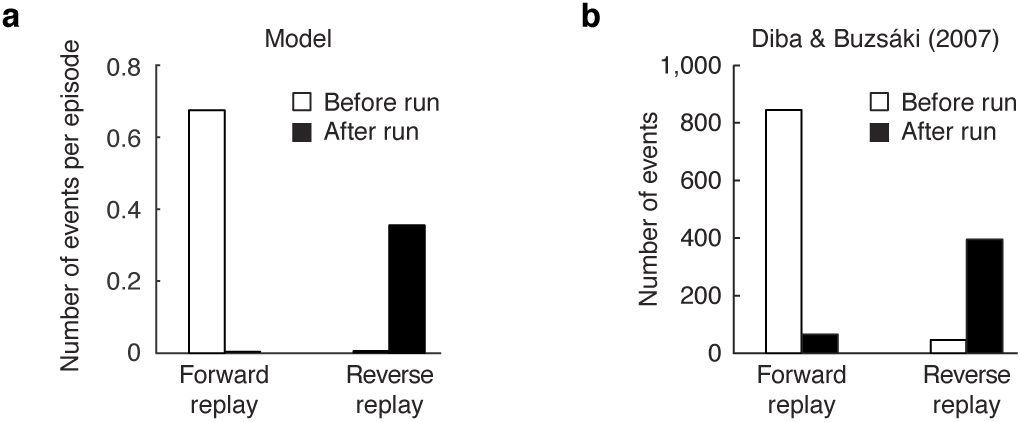
Forward and reverse sequences happen at different times and are modulated asymmetrically by reward. *(a)* Forward sequences tend to take place before the onset of a run while reverse sequences tend to take place after the completion of a run, upon receipt of reward. *(b)* Data from Diba & Buzsáki (2007), their Fig. 1C.

### 2.3 Statistics of replayed locations: current position, goals, and paths

In addition to directionality, the theory predicts which particular forward or backward routes should be considered, which ultimately determines the locations of behavioral change. In general, replay should be biased toward relevant locations in the environment such as the agent’s current position (due to high need) and reward sites (due to high gain). These general biases arise from the average over many situations of its more specific experience-and situation-dependent replay, which is patterned due to the influence of particular locations like reward sites on both the need and gain terms.

In principle, forward and reverse sequences can begin anywhere along the linear track. Yet, in our simulations most significant events start in locations at or immediately behind the agent and extend thereafter in either direction to propagate value (Fig. 4a). Empirical results on the linear track support this prediction: hippocampal events have an overall tendency to reflect a path that begins at the animal’s current location^21,23,24^, a phenomenon termed “initiation bias” (Fig. 4b).

**Figure 4:**
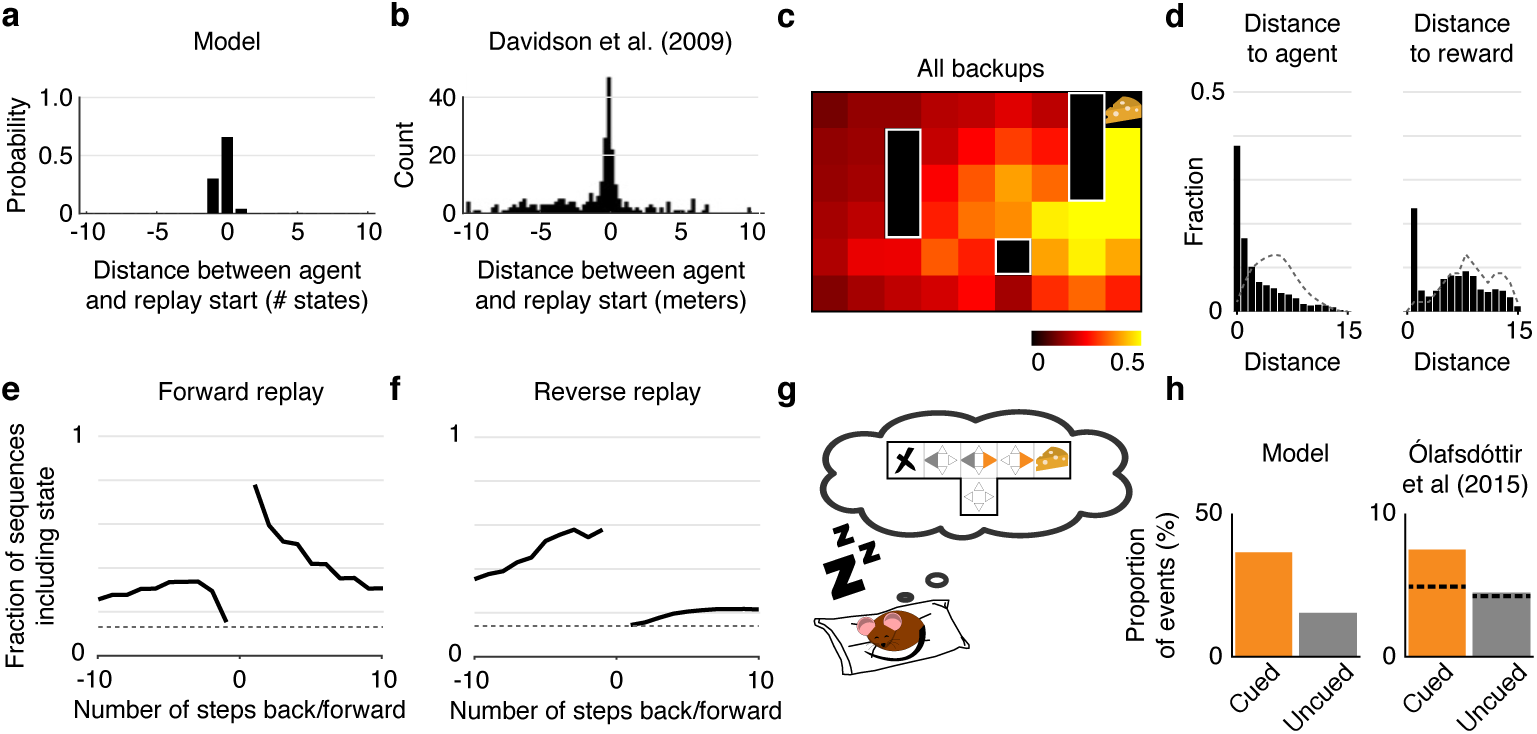
Replay over-represents agent and reward locations and predicts subsequent and past behavior. *(a)* Distribution of start locations of significant replay events relative to the agent’s position and heading on the linear track. Negative distances indicate that the replayed trajectory starts behind the agent. Most significant replay events in the linear track start at or immediately behind the agent’s location. *(b)* Data from Davidson et al (2009) showing the distribution of start locations of replay trajectories relative to the animal’s position and heading on the track. *(c)* Activation probability across all backups within an episode. Colors represent the probability of a backup happening at each location within a given episode. Notice that backups are more likely to occur in locations near the reward. *(d)* Probability that a given backup happens at various distances from the agent (*left*) and from the reward (*right*) in the open field. Dotted lines represent chance levels. Notice that backups are substantially more likely to happen near the agent and/or near the reward than chance. *(e,f)* How forward and reverse replay predict future and previous steps in the open field. The lines indicate the probability that the first 5 backups of any significant forward or reverse sequence contains the state the agent will/have occupied a given number of steps in the future/past. Dotted lines represent chance levels. Notice that forward replay is more likely to represent future states than past states, while the opposite is true for reverse replay. *(g)* We simulated an agent in an offline setting (e.g. sleep) after exploring a T-maze and receiving a reward on the right (cued) arm. *(h) Left*: The proportion of backups corresponding to actions leading to the cued arm (orange) is much greater than the proportion of backups corresponding to actions leading to the uncued arm (gray). *Right*: Data from Ólafsdóttir et al (2015) showing the proportion of spiking events categorized as “preplay” events for the cued and uncued arms. The dashed line indicates the proportion of events expected by chance.

Sequences that start at the animal’s current location can, nonetheless, extend in any direction, in particular in open field environments where trajectories are less constrained by the environment. Yet, gain and need in the model both favor important locations like goals or reward sites, and empirically, sequential replay in open environments is also biased toward these locations^22,43^. We simulated navigation in an open field environment (Fig. 1c, *left*) and examined these content biases by calculating the *activation probability* for each spatial location in the open field environment (the probability of a backup happening at each state within an episode). Visualized over space (Fig. 4c), backups tended to concentrate near the reward (goal) locations, in line with rodent recordings^22,44^. Quantified as a function of distance (Fig. 4d), backups were again more likely than chance to happen near the reward and/or the agent (the latter result replicating in the open field the initiation bias from Fig. 4a^22,45^.

Results like these have been taken to reflect replay’s involvement in planning future routes, and indeed the bias toward locations near the goal was seen even when forward replay was considered separately (Fig. S1), which cannot simply reflect initiation bias because starting locations were randomized in our simulations. Interestingly, locations corresponding to the final turn toward the reward were emphasized even more than locations nearer the reward itself, a consequence of the gain term being higher where there is a greater effect on behavior. The over-representation of turning points is a consequence of the barriers in the simulated environment, which produce sharp turning points, and is consistent with empirical reports that reactivated place fields congregate around relevant cues^46^.

The hypothesized involvement of replay (both forward and reverse) in evaluating potential routes can also be assessed by comparing replayed trajectories to recent or future paths. In the model, these tend to coincide, both because backups tend to occur in locations favored by the need term, and additionally, in the case of forward trajectories, by the definition of valid *n*-step sampling, which measures rewards expected along the predicted future trajectory of the agent. However, the correspondence is not perfect; in fact backups can sometimes construct trajectories not previously traversed continuously by the agent^25^. (Although our model as implemented only replays individual transitions that have previously been made in real experience, the same framework would work equally with transitions whose availability and potential future need can be inferred, as by vision.) We measured the probability that the first 5 backups of a forward or reverse event would include locations visited by the agent in the future or past. In the open field, our simulations revealed that forward replay has a higher probability than chance of including states visited a few steps in the future, and a probability only slightly higher than chance to include states visited in the past (Fig. 4e). In contrast, reverse replay has a higher probability than chance of including states visited in the past, and a probability only slightly higher than chance to include states visited in the future (Fig. 4f). That replayed trajectories tend to correspond to the specific trajectories followed by the agent in either the past (reverse replay) or future (forward replay) is again in line with rodent studies^22,46^.

Lastly, we address the case of remote replay, where sequences correspond to spatial locations away from the animal^24^ or remote environments altogether^28^. In particular, even during sleep — where the content of replay rarely corresponds to the location where the animal is sleeping — replay tends to represent rewarding areas of the environment in comparison to similar but unrewarding areas^44^. In our model, biases in reactivation during rest can again be understood in terms of the same need-and gain-based prioritization (with need defined as expected future occupancy subsequently). We tested these predictions of sleep replay by simulating a T-maze with a reward placed at the end of one of the two arms (Fig. 4g), with the agent absent from the environment (see Methods). The proportion of backups corresponding to actions leading to the cued (rewarded) arm was much greater than the proportion of backups corresponding to actions leading to the uncued (unrewarded) arm (Fig. 4h), reproducing equivalent empirical results^44^.

### 2.4 Asymmetric effect of prediction errors

We have shown that prioritized memory access for action evaluation applied in different conditions may give rise to forward and reverse sequences. However, our claim that both forward and reverse replay may arise from the same prioritized operation may seem at odds with the general view that forward and reverse sequences have distinct functions (e.g., planning and learning, respectively^21,26^). One piece of evidence that has been argued to support this distinction is the observation that reverse and forward replay have different sensitivities to reward context: in rodents navigating a linear track, the rate of reverse replay is increased when the animal encounters an increased reward, and decreased when the animal encounters a decreased reward. In contrast, the rate of forward replay is similar despite increases or decreases in reward^26,46^.

Our hypothesis is instead that planning and learning are better understood as different variants of the same operation, i.e. using backups (in different orders) to propagate reward information over space and time. In our model, asymmetric effects of increases vs. decreases in reward are a hallmark of the gain term, arising from its definition in terms of policy change (Fig. 1e,f), and distinguishing our prioritization hypothesis from others that simply trigger update on any surprise^36–38^).

Because gain is accrued when an update changes the agent’s choices toward a better one, it depends both on whether the news is good or bad, and also what alternative actions are available (Fig. 1e,f). Fig. 5a,b demonstrates this predicted interaction by plotting gain for different types of feedback about the action previously believed to be better (Fig. 5a) or worse (Fig. 5b) in a two-action situation. The basic pattern is that gain is small for learning that the better action is even better, but large for learning it is actually much worse than the alternative (since it teaches the agent the value of switching actions). The pattern reverses for updates to the worse action’s value: learning that it is better than the alternative carries gain (since it can now be chosen) but learning it is even worse than previously believed is not helpful.

**Figure 5:**
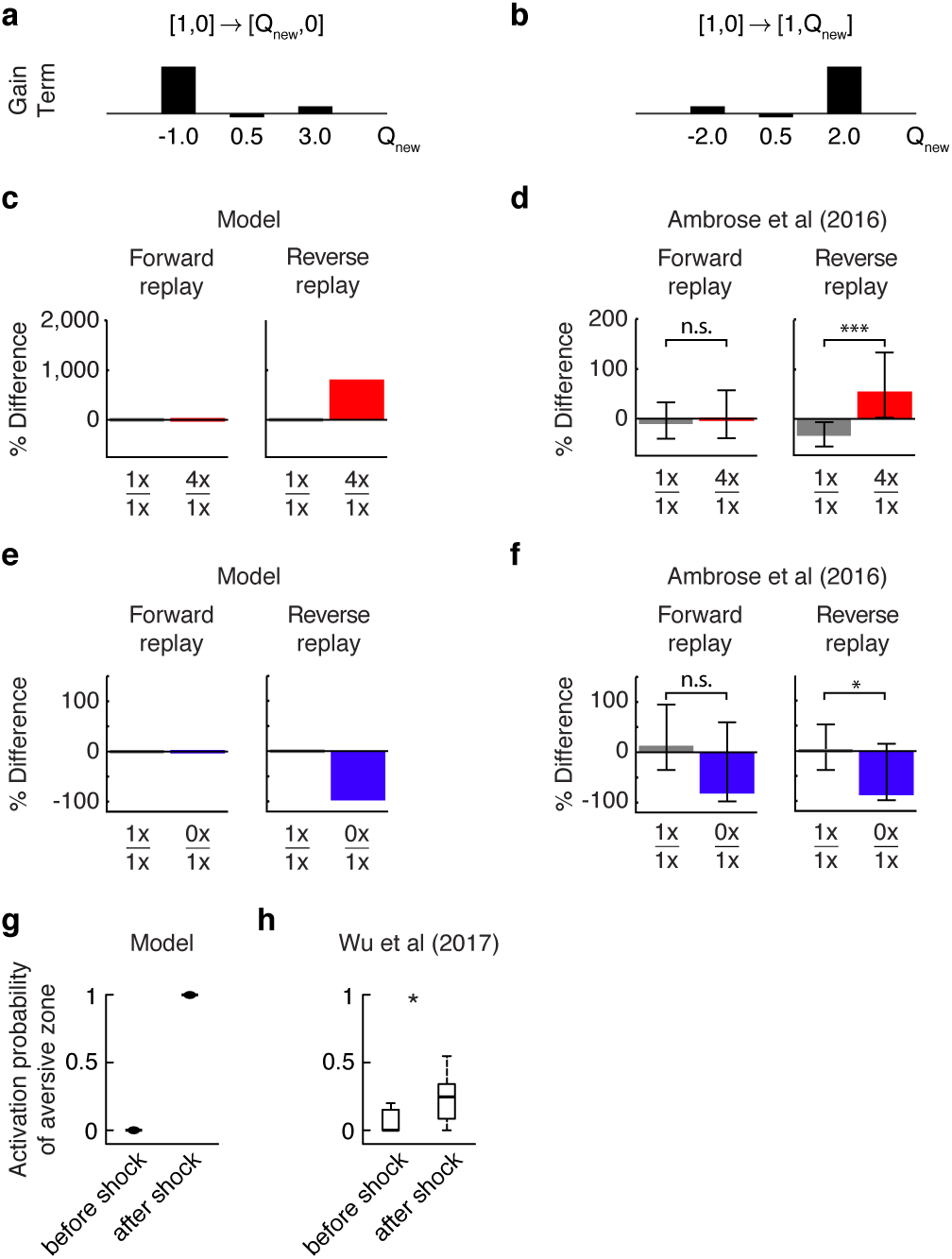
Forward and reverse sequences happen at different times and are modulated asymmetrically by reward. *(a)* Gain term for an example case where two actions are available and the agent learns a new value (*Q_new_*) for the best action. *(b)* Gain term for an example case where two actions are available and the agent learns a new value (*Q_new_*) for the worst action. *(c)* We simulated a task where, in half of the episodes, the reward received was 4x larger than baseline. *Left*: The number of forward events was approximately equal in every lap both when the rewards were equal, as well as when the rewards were 4x larger. *Right*: In contrast, the number of reverse events was approximately equal when the rewards were equal, but much larger upon receiving a larger reward in the unequal reward condition. *(d)* Data from Ambrose et al (2016), their Fig. 3E,H. *(e)* We simulated a task where, in half of the episodes, the reward received was zero. *Left*: The number of forward events was approximately equal in every lap both when the rewards were equal, as well as when the rewards were removed. *Right*: In contrast, the number of reverse events was approximately equal when the rewards were equal, but almost completely abolished upon receiving no reward in the unequal reward condition. *(f)* Data from Ambrose et al (2016), their Fig. 5C,F; note that the effect of reward for forward replay (left) is not significant. *(g)* Activation probability at the end of a linear track during random exploration without rewards or punishments (*left*) and after shock delivery at the end of a track (*right*). *(h)* Data from Wu et al (2017) (their Fig. 3e) showing the activation probability during population burst events of cells with place fields at a light zone before shock delivery (*left*) and similarly for cells with place fields at a shock zone after shock delivery (*right*).

When learning about the better action, there is also an additional, subtler asymmetry in gain between good and slightly bad news, due to changes in the *degree* of preference for the better action depending on its relative advantage (softmax exploration). Because learning the better action is even better results in an updated policy in which the agent is more likely than previously to take it, this does carry positive gain. Conversely, if the reward is found to be smaller than the corresponding action value, but still equal to or better than the alternative, gain will be zero or negative, reflecting an updated policy where the agent is less likely to collect whatever reward is remaining and more likely to take an inferior exploratory action (Fig. 5a,b). Altogether, when the best action is taken, novel reward information will tend to be be propagated backwards only for increases in reward, or decreases worse than any alternative action. Notice that these effects are predicted to arise only for reverse replay occurring at the end of a run, when the gain term is large and, therefore, dominates the utility of the backup.

There are empirical data supporting several aspects of these predictions. Consider the second predicted asymmetry: between better or worse news about an action that is nevertheless still better than any alternative. We investigated the differential response of these two types of replay in these situations by simulating two conditions on a linear track task similar to that studied by Ambrose et al. (2016)^26^: (i) an *increased reward* condition where the reward encountered by the agent was four times larger in half of the episodes, and (ii) a *decreased reward* condition where the reward encountered by the agent was zero in half of the episodes. The number of forward events was approximately equal in the increased reward setting regardless of the reward received. In contrast, the number of reverse events was much larger upon receiving a larger reward than upon receiving a conventional reward (Fig. 5c,d). This effect was driven both by an increase in the rate of reverse replay for larger rewards, and a decrease for conventional (1x) rewards (Fig. S2), as observed experimentally^26^. In the decreased reward setting, the number of forward events was also approximately equal in every lap regardless of the reward received. In contrast, the number of reverse events was much smaller upon receiving no reward than upon receiving a conventional reward (Fig. 5e,f). This effect was driven both by a decrease in the rate of reverse replay when the reward was 0, and an increase when the reward was conventional (1x) (Fig. S2), again replicating empirical findings^26^.

Notice that, in principle, these asymmetries will tend to slow the updating of values following certain types of news, biasing Q-values up or down in different situation. Over time, however, conventional *Q*-learning over real experience, will tend to progressively correct these biases. Also note that Ambrose et al (2016)’s tasks have an additional feature that, in our model, also disfavors propagating information about zero rewards: the animal must visit the end of the track before collecting a reward on the other end. In this setting, propagating information about a reward of 0 is prejudicial (negative gain), as it reduces the probability of the animal reaching the end of the track, delaying the collection of the other (1x) reward.

The second asymmetry predicted by our model concerns the usefulness of finding out the previously best action is *worse* than alternatives. For this, consider a scenario in which a negative reward (e.g., an electric shock) is encountered at the end of the track. In this case, discovering this shock has a positive gain (Fig. 5a), as it enables a favorable behavioral change: omitting the action altogether. A crucial prediction of the model, therefore, is that propagating a negative prediction error is unhelpful if the alternative actions would still be disfavored, but advantageous if the alternative actions become preferred. Notably, staying still or moving backwards for no outcome is better than moving toward a frankly aversive shock zone. Indeed, in our simulations, there is a unit probability of a backup happening at the shock zone after shock delivery (to propagate this information and prevent the agent’s return to the shock zone), in contrast to a negligible probability of a backup happening in the same area prior to shock delivery (Fig. 5g). This prediction has also been confirmed recently: in a conditioned place avoidance task, replay sequences were observed extending from the animal’s position toward the end of a linear track previously paired with a shock, despite the fact that the animals did not then enter the shock zone^47^ (Fig. 5h). These results not only provide direct support to our theory’s notion of gain, but also illustrate how the notion of planning embodied by our model differs from a narrower, colloquial sense of planning: Evaluating candidate actions by simulation (as in our model) does not just anticipate paths to goals, it can also help agents figure out what *not* to do.

### 2.5 Effects of familiarity and specific experiences

As a learning model, our theory also predicts characteristic effects of experience on the prevalence and location of replay. In particular, change in the need vs. gain terms predicts countervailing effects of task experience. As a task becomes well learned, prediction errors decrease, policies stabilize, and the gain expected due to replay decreases, causing a reduction in significant replay events (in the model, we encourage continuous learning by adding a small amount of noise to the rewards and assessing zero gain at a small nonzero minimal value, meant to capture an assumption of persistent uncertainty due to the possibility of environmental change). At the same time, as behavior crystallizes, the need term becomes more focused along the particular routes learned by the agent (e.g., compare Fig. 1g, *top* and *bottom*). This predicts that, conditional on replay occurring, particular states are increasingly likely to participate.

The balance between these countervailing effects may help to explain apparent inconsistencies in the replay literature, as both increases and decreases in replay have been reported, albeit using a range of different dependent measures and designs. Indeed, existing reports on the effect of familiarity on the incidence of replay have been difficult to reconcile^45,46,48–50^. While some studies report an increase in reactivation with time spent in the environment^45,48^, several others report a decrease with familiarity^21,23,46,49^. Specifically, the more time an animal spends between two place fields, the more the corresponding place cell pair is reactivated during sleep (consistent with focusing of need on these states^48^). In contrast, replay is more easily observed in novel than in familiar tracks (consistent with a decrease in gain overall^23^), and the average activation probability is highest in novel environment^49^. It has been suggested that replay tends to increase within session with exposure, but decrease across sessions as the animal becomes familiar with a novel environment^50^. This may reflect the additional effect of experience vs. computation on learning in our model. In particular, both need (favoring focused replay) and gain (opposing overall replay) are affected by actual experience in an environment, but only gain is affected by replay (e.g. during rest between sessions). This is because only experience can teach an agent about the situations it is likely to encounter (i.e. need), but value learning from replayed experience reduces subsequent gain.

We examined the effect of familiarity and specific experience on replay by calculating the number of significant replay events as a function of experience (episode number). In line with previous reports^23^, we observed that the number of significant events is highest during the first trials and decays steadily with experience, going from about 2 to about 0.8 significant events per trial. This effect was due to a decrease in both forward replay (going from about 1.5 to 0.5 events per trial), and reverse replay (going from about 0.5 to 0.3 events per trial). A similar effect was also observed in terms of activation probability: the average probability of a random state being part of a replay event. Similarly, activation probability decayed steadily with experience (Fig. 6a), in line with empirical findings (Fig. 6b)^49^, and this decay was observed for both forward and reverse sequences (Fig. 6a, insets). In contrast, we observed that the probability of an event including a specific state increased with number of visits (Fig. 6c), also in line with previous reports (Fig. 6d)^48^. Again, these two effects reflect the effect of experience on the two terms governing priority: while the gain term decreases with exposure (with the gradual reduction in prediction errors), the need term increases as the agent’s trajectory becomes more predictable.

**Figure 6:**
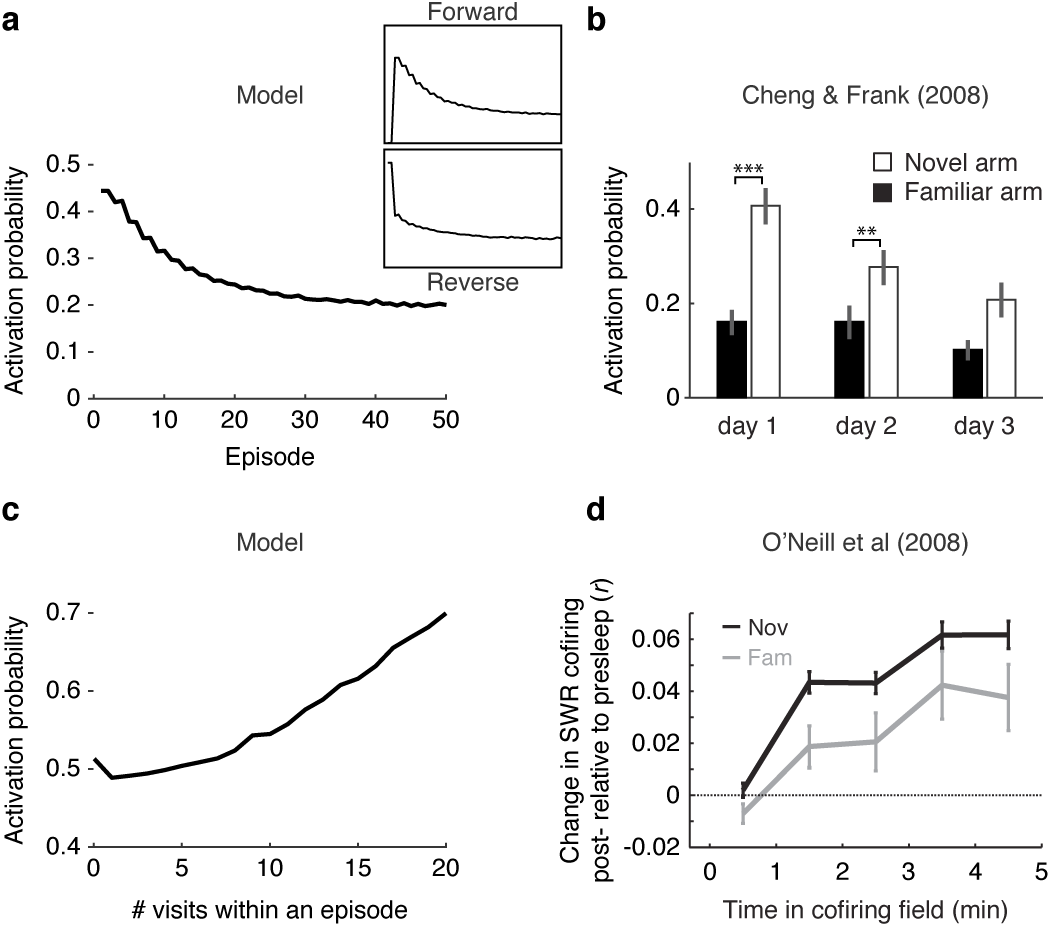
Replay frequency decays with familiarity and increases with experience. *(a)* In the linear track, the probability that significant replay events include a state in the linear track decays across episodes, peaking when the environment is novel. Insets show that the number of both forward (*top*) and reverse (*bottom*) replay events decay with experience. *(b)* Data from Cheng Frank (2008) showing the activation probability per high-frequency event. Error bars represent standard errors and symbols indicate results of rank-sum test (** *p* < 0.01, *** *p* < 0.001). *(c)* Probability that significant replay events include a state in the linear track as a function of the number of visits in an episode. Analogously to the effect reported in Fig. 1g, driven by the need term, the probability of a state being replayed increases with experience in that state. *(d)* Data from O’Neil et al (2008) showing that that the more cell pairs that fire together during exploration (time in cofiring field), the larger is the increase in probability that these cell pairs fire together during sleep SWRs.

### 2.6 Effect of replay on choice behavior

The preceding simulations demonstrate that a wide range of properties of place cell replay can be predicted from first principles, under the hypothesis that the common goal of replay is to drive reinforcement learning and planning involving the reactivated locations. This hypothesis also makes a complementary set of predictions about choice behavior, i.e. that replay will be causally involved in learning which actions to take at the replayed states. Such behavioral effects are most characteristically expected for acquiring tasks (like revaluation and shortcut tasks) that require agents to integrate associations learned separately, or to infer the value of novel actions. This is because this class of tasks cannot be solved by alternative learning mechanisms in the brain (such as model-free TD learning, associated with dopamine and striatum) and exercise the more unique ability of nonlocal replay to compose novel trajectories from separate experiences^3^.

Indeed, hippocampal replay can follow novel paths or shortcuts never traversed by an animal^25^, and in one report^44^, activation of a path not yet explored (because it was initially observed behind glass) was followed by rats subsequently being able to choose that path, correctly, over another, consistent with the planning hypothesis. While our simulations required the explicit experience with each action for it to be replayed, it is reasonable to assume that replay is already possible with mentally assembled experienced from other senses such as vision, so these findings are generally consistent with our theory. In the open field, forward hippocampal replay predicts future paths, importantly even when the goal location is novel^22^. Finally, causally blocking sharp wave ripples has a selective effect on learning and performance of a spatial working memory task^31^; although the specifics of that task would require elaboration of our model to simulate (mainly because it turns on a history-dependent response rule whereas we consider strictly Markovian tasks), it provides a clear demonstration that awake replay is required for associating events over space and time^31^. Overall, though, the place cell literature has tended not to focus on connecting hippocampal activity to learning and decision behavior, and a major area for future research suggested by our theory is to combine tasks more specifically diagnostic of model-based planning and reinforcement learning with the monitoring and manipulation of nonlocal replay.

A second feature of the current theory is that it emphasizes that a number of different patterns of replay (forward, reverse, and offline; Fig. 1a) can all equally be used to solve the sorts of integrative decision tasks that have largely been assumed to reflect forward “model-based” planning at the time of choice. Indeed, forward planning of this sort may be subserved by “preplay”^20,22^, but in the current theory, this is just one case of a more general mechanism, and equivalent computations can also arise from memories activated in different patterns at other times. These also include reverse replay that allows connecting an experienced outcome with potential predecessor actions, and nonlocal replay composing sequences of experiences during rest. Although these possibilities have not been examined in hippocampal spatial literature, work with humans using non-spatial versions of revaluation tasks (and activity of category-specific regions of visual cortex to index state reinstatement) verifies that not just forward replay^5^ but also reverse replay^13^ and nonlocal replay during rest^15^ all predict the ability of subjects to solve these tasks. The present theory’s account of which replay events are prioritized might provide a basis for investigating why different studies and task variants tend to evoke one or the other of these solution strategies.

## 3 Discussion

In light of so much experience accumulated in a lifetime, which memories should one access and when to allow for the most rewarding future decisions? We offer a rational account for the prioritization of memory access operations framed in terms of action evaluation through Bellman backups. We propose that the various nonlocal place cell phenomena in the hippocampus reflect different instances of a single evaluation operation, and that differences in the utility of these operations can account for the heterogeneity of circumstances in which they happen. This utility, derived from first principles, amounts to the product of a gain and a need term. Simulations of the model reproduced qualitatively a wide range of results reported in the hippocampal replay literature over the course of the previous decade without the need for parameter fitting.

This theory draws new, specific connections between research on the hippocampal substrates of spatial navigation, and research on learning and decision making, with implications for both areas. It has long been recognized that place cell activity (including forward and reverse replay) likely supports learning and decision making^20,23^; the present research renders these ideas experimentally testable by offering a specific hypothesis about what the brain learns from any particular replay event. This immediately suggests experiments combining trial-by-trial reward learning (of the sort often studied in the decision literature) with place cell monitoring, to test the the predicted relationship between individual replay events and subsequent choices. The strongest test would use tasks (like sensory preconditioning or multi-step sequential decision tasks) where relevant quantities cannot be directly learned “model-free” over experienced trajectories but instead exercise the ability of replay to compose novel sequences^3,15,51^. Upstream of this behavioral function, the theory also makes new testable predictions about place cells themselves, by articulating quantitative experience-and circumstance-dependent criteria for which locations will be replayed.

The hippocampal literature has tended to envision that replay serves disjoint functions in different circumstances, including learning^23^, planning^20–22,52^, spatial memory retrieval^31^, and systems consolidation^30,32^. By focusing on a specific, quantitative operation (long-run value computation), we sharpen these suggestions and expose their deeper relationship to one another. In RL, learning amounts to propagating long-run value information from one state to adjacent ones to perform temporal credit assignment, with forward “planning” as traditionally conceived being one special case. This perspective unifies the proposed role of forward replay in planning with that of reverse replay in learning (both linking recently experienced sequences to their outcome^23^), and suggests a similar role for analogous nonlocal computations, e.g. during sleep. Though serving a common goal, these different patterns of replay are most appropriate in different circumstances; this explains observations of differential regulation (such as asymmetric effects of prediction errors on forward vs. reverse replay), which have otherwise been taken as evidence for distinct functions^26^. As for consolidation, the perspective that replay drives estimation of long-run values echoes other work on systems consolidation^32,53^ in viewing consolidation not merely as strengthening existing memories, but more actively computing new summaries from the replayed content. Also as with other systems consolidation theories, the resulting computed quantities (here, action values) are widely believed to be stored elsewhere in the brain (here, likely cortico-striatal synapses), and the fuller neural processes of replay presumably involve coordinated evoked activity throughout the brain, especially including value prediction and learning in the mesolimbic and nigrostriatal reward networks^41,42^.

Relatedly, while we have hypothesized a specific role for replay in computing long-run action values — and although it is striking that this consideration alone suffices to explain so many regularities of place cell replay — we do not view this function as exclusive of other computations over replayed experiences^32,53^. One interesting variant of our theory is that replay can be used to learn a long-run transition model of the particular locations and outcomes one expects to visit following some action — instead of, as in our theory, the long-run reward consequences alone. Such a long-run outcome representation, known as the successor representation (SR), can serve as an intermediate representation for computing action values^54^, a sort of temporally extended cognitive map. The SR has recently been proposed to be learned within hippocampal recurrents^55^ and to explain aspects of human choice behavior^56^. It can also be updated using replayed experience (“SR-DYNA”^57^) analogous to how we learn reward values here, connecting learning from replay more directly with a type of cognitive map building^25^. Our ideas carry directly over to this case: In fact, our prioritization computations remain unchanged if replay updates an SR instead of action values; this is because an SR update step (also based on the Bellman equation) has exactly the same utility (under our myopic approximation) as the corresponding Bellman backup for action values.

From the perspective of decision neuroscience, a key driver of recent progress is the recognition that the details how decision variables are computed — specifically, whether an action’s consequences are considered — govern what choices are made. Notably, the view that the brain contains separate systems for “model-based” vs. “model-free” value computation (which differ in whether or not they recompute values via planning at decision time) offers an influential computational reframing of issues such as habits and compulsion. Yet a realistic “model-based” system must necessarily be selective as to which of many branches are considered^12,58,59^, and dysfunction in such selection may extend the reach of these mechanisms to explain symptoms involving biased (e.g., abnormal salience or attention in both compulsive and mood disorders; craving) and abnormal patterns of thought (e.g., rumination, hallucination). The current theory goes beyond just prioritizing planning about the immediate future, to also consider value computation at nonlocal states not immediately implicated in decision, e.g. “offline” replay during sleep or rest^29,34,60^. This systematizes several instances by which tasks typically thought to index “model-based” planning at choice time are apparently solved by computations occurring earlier^13–15^ and suggests links between these phenomena and different patterns of replay. Finally, by recasting planning as learning from remembered experience, the theory envisions that the value learning stage of it might be subserved via the same dopaminergic error-driven learning operation long thought to support model-free learning from direct experience. This more convergent picture of the substrates of these two sorts of learning would explain results (puzzling on a separate systems view) that dopaminergic activity is both informed by^61,62^ and supports^63–65^ model-based evaluation.

The AI literature suggests one other candidate hypothesis for the prioritization of backups, known as Prioritized Sweeping (PS)^36,37^ and its recent variants used in Atari game playing^38^. The idea is that large prediction errors (whether negative or positive) should drive backup to propagate the unexpected information to predecessor states. Our approach adds the need term (focusing backups on states likely to be visited again). Also, in the gain term, it takes into account the effect of a backup on an agent’s policy, ensuring that propagating information with no behavioral consequences has no value. Data support both of these features of our model over PS: the gain term (unlike PS) predicts asymmetric effects of positive and negative prediction errors^26^ (Fig. 5c-f). Because of the need term, our model can also produce searches forward from the current state, in addition to PS’s largely backward propagation of error. The need term has a second effect, which is to channel sequential activity along recently or frequently observed trajectories. This may help to explain why nonlocal place cell activity follows extended sequences even though a straightforward error propagation is often more breadth-first^36,37^.

The need term also bears close resemblance to the concept of *need probability* from rational models of human memory^66^ — the probability that an item needs to be retrieved from memory because of its relevance to the current situation. Indeed, although we have framed our theory in terms of memory access, our use of a deterministic, static task moots important distinctions between different sorts of memory, such as episodic and semantic. In particular, replay-based methods can learn equivalently either from remembered experiences (e.g., episodic memories of particular trajectories), or from simulated experiences (e.g., trajectories composed by first learning a semantic map or model of the task, then generating experiences from it, as in model-based RL), blurring the distinction between model-based learning and model-free learning from stored samples^67^. In the current setting, prioritizing over experiences, locations, and maps all amount to the same thing, since due to the nature of the task, any episode of going (for instance) north from a particular location is identical to any other. An important goal of future work will be to tease apart the role of episodic vs. semantic knowledge in computing action values, and understand their relative prioritization^68,69^.

There are a number of other limitations to the model, many of which are also opportunities for future work. Though we have used a spatial framing due to the links with hippocampal replay, our theory is formalized generally over states and actions and can be applied beyond navigation to other sequential tasks. However, we omitted many model features to construct the simplest instantiation that most clearly exposes the key intuition behind the theory: the interplay between gain and need and their respective roles driving reverse and forward replay. For instance, we restricted our simulations to two very simple environments (a linear track and an open field). While grid-world simulations are obviously poor approximations of real navigation (animals clearly do not start from a random walk strategy nor treat adjacent states independently), our goal was not to produce an accurate model of rodent behavior but to expose the normative principles governing memory replay. Relatedly, we assumed a stationary and deterministic environment that can be learned by the agent without uncertainty — and accordingly omitted stochasticity and uncertainty from the model also. Yet, a full account of prioritized deliberation must surely account for uncertainty about the action values and its sources in stochasticity and nonstationarity. This will require, in future, re-introducing these features from previous accounts of online deliberation^7,8^; with these features restored, the current theory will inherit its predecessors’ successful account of phenomena of habits, such as how they arise with overtraining.

This also relates to perhaps the most important limitation of our work: to investigate the decision theoretic considerations governing replay, we define priority in the abstract, and do not offer a mechanism or process-level recipe for how the brain would realistically compute it. Although the need term is straightforward (it corresponds to the SR^54^, which the brain has already been proposed to track for other reasons^55,56^), the calculation of gain, as we define it, requires that the agent knows the effect of the backup on its policy prior to deciding whether to perform the backup. We use this admittedly unrealistic decision rule to investigate the characteristics of efficient backup, but a process-level model will require heuristics or approximations to the gain term; here again previous work on deliberation under uncertainty suggests a candidate approximation, called the myopic value of perfect information^8^.

To highlight the role of sequencing computations and minimize assumptions about the nature of the replayed events, we have also constructed the theory at a single spatial and temporal scale, focusing on a single Bellman backup as the elementary unit of computation. We build both forward and reverse replay trajectories recursively, step by step, with value information propagating along the entire trajectory. Of course, research in both hippocampus and decision making (separately) stresses the multi-scale nature of task representations. A fuller account of learning, planning, and prediction would include temporally extended actions (“options”)^59,70,71^ or similar long-scale state predictions^54,72^. In this case, the principles of prioritization would carry over directly, but the elemental replay events being considered would be structured as extended trajectories and treated as ballistic events, rather than individual locations.

## 4 Methods

### 4.1 Model description

The framework of reinforcement learning^1^ formalizes how an agent interacting with an environment through a sequence of states should select its actions so as to maximize some notion of cumulative reward. The agent’s policy *π* assigns a probability *π*(*a*|*s*) to each action 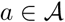 in state 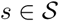. Upon executing an action *A_t_* at time *t*, the agent transitions from state *S_t_* to state *S_t_*_+1_ and receives a reward *R_t_*. The goal of the agent is to learn a policy that maximizes the discounted return *G_t_* following time *t* defined as:

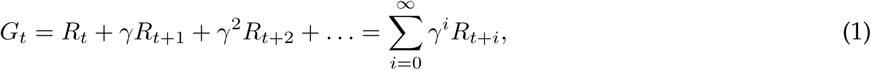
where *γ* ∈ (0, 1] is the *discount factor* that determines the present value of future rewards.

The expected return obtained upon performing action *a* in state *s* and subsequently following policy *π* is denoted *q_π_*(*s, a*) and is given by:

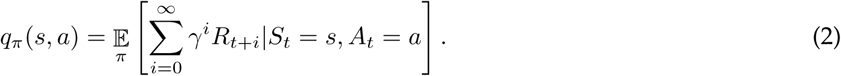

The policy that maximizes the expected return is the *optimal policy* and denoted *q*_*_. Following *Q*-learning^73^, the agent can learn an action-value function *Q* that approximates *q*_*_ through iteratively performing Bellman backups:

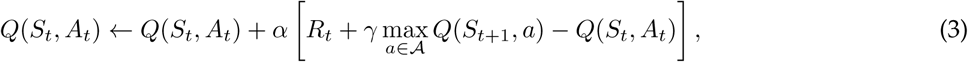
where *α* ∈ [0, 1] is a learning rate parameter. Bellman backups are performed automatically after each transition in real experience and may also be performed nonlocally during simulated experience, as in the DYNA architecture^34^.

The following framework provides a rational account for prioritizing Bellman backups according to the improvement in cumulative reward expected to result. Let the agent be in state *S_t_* = *s* at time *t*. We represent an experience *e_k_* by the 4-tuple *e_k_* = (*s_k_,a_k_,r_k_,s′_k_*), and we consider that accessing experience *e_k_* amounts to a Bellman backup which updates *Q*(*s_k_,a_k_*) with the target value 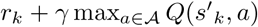. We also denote by *π_old_* the current (old) policy, prior to executing the backup, and *π_new_* the resulting (new) policy after the backup.

The utility of accessing experience *e_k_* to update the value of *Q*(*s_k_,a_k_*), or Expected Value of Backup, is denoted by *EV B*(*s_k_,a_k_*) and is defined as:

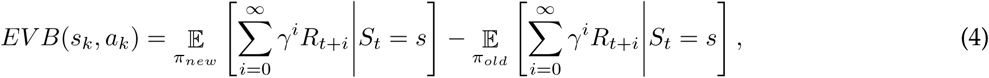
i.e., EVB is the improvement in expected return due to a policy change. A key point about this definition is that although it sums rewards over all future timesteps, it can be rewritten in terms of a sum over expected visits to the updated state *s_k_* (the full derivation is given below.) This is because accessing *e_k_* can only affect the policy in state *s_k_* (i.e., *π_new_* and *π_old_* differs only in state *s_k_*); and we can then separately consider the gain accrued each time the agent visits that state *s_k_*, and the expected number of times *s_k_* will be visited. In other words, by conditioning *EV B*(*s_k_,a_k_*) on *S_t_* = *s_k_*, this expression can be separated into the product of two terms: *EV B*(*s_k_,a_k_*)= *Gain*(*s_k_,a_k_*) × *Need*(*s_k_*).

#### 4.1.1 Gain term

The gain term quantifies the expected improvement in return accrued at the target state, *s_k_*:

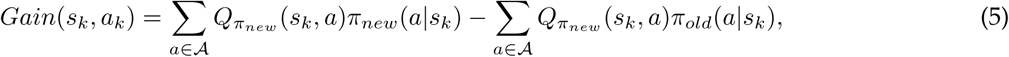
where *π_new_*(*a*|*s_k_*) represents the probability of selecting action *a* in state *s_k_ after* the Bellman backup, and *π_old_*(*a*|*s_k_*) is the same quantity *before* the Bellman backup.

#### 4.1.2 Need term

The need term measures the discounted number of times the agent is expected to visit the target state, a proxy for the current relevance of each state:

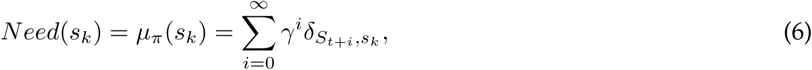
where *δ*_·,·_ is the Kronecker delta function. Notice that, for *γ* =1, the need term is the exact count of how many visits to state *s_k_* are expected in the future, starting from current state *S_t_* = *s*.

The need term can be estimated by the Successor Representation^54^, which can be learned directly by the agent or computed from a model. Here, we assume that the agent learns a state-state transition probability model 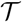 for the purpose of computing the need term. The need term is thus obtained directly from the *n*-th row of the SR matrix, 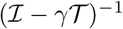, where *n* is the index of the agent’s current state *S_t_*.

An alternative option is to use the stationary distribution of the MDP, which estimates the asymptotic fraction of time spent in each state (i.e., after convergence). This formulation is particularly useful when the transition probability from the agent’s current state is unavailable (e.g., during sleep).

### 4.2 Simulation details

We simulated two “grid-world” environments (Fig. 1c) where an agent could move in any of the four cardinal directions – i.e. 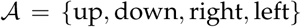. At each state, the agent selects an action according to a softmax decision rule over the estimated *Q*-values, 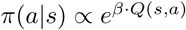, where *β* is the inverse temperature parameter which sets the balance between exploration versus exploitation. In our simulations, *β* = 5. Upon selecting action *A_t_* = *a* in state *S_t_* = *s*, the agent observes a reward *R_t_* = *r* and is transported to an adjacent state *S_t_*_+1_ = *s*′. The value of *Q*(*s, a*) is then updated according to (3) using *α* = 1.0 and *γ* = 0.9. We used a learning rate of *α* = 1 due to it being both maximally simple and optimal when the world’s dynamics are deterministic.

The first environment — a linear track (Fig. 1c, *right*) —, was simulated as two disjoint 1 × 10 segments. (The motivation for this was for the state space to differentiate both location and direction of travel, as do hippocampal place cells in this sort of environment; this also clearly disambiguates forward from reverse replay.) The agent started in location (1, 1) of the first segment. Upon reaching the state (1, 10), the agent received a unit reward with Gaussian noise added with standard deviation of *σ* = 0.1 and was transported to state (1, 10) of the second segment. Upon reaching state (1, 1) in the second segment, the agent received a new unit reward (plus independent Gaussian noise with *σ* = 0.1) and was transported back to state (1, 1) of the first segment. Each simulation comprised of 50 episodes (i.e. sequence of steps from starting location to reward). The second environment was a 6 × 9 field with obstacles (Fig. 1c, *left*), with a unit reward (*σ* = 0.1) placed at coordinate (1, 9). Each simulation comprised of 50 episodes with the start location randomized at each episode.

Our theory assumes that the memory access leads to more accurate *Q*-values. Improved estimates of action values can be obtained from samples of experience in which that action is used. We consider the need and gain for activating all one-step experiences *e_k_* = (*s_k_,a_k_,r_k_,s′_k_*); these generally correspond to one-step updates given by (3). In addition, we consider one special case: if the target state action (*s_k_,a_k_*) is an optimal continuation of the sequence replayed immediately previously (i.e. if *s_k_* was the end state considered previously, and *a_k_* is the optimal action there), then this replay can extend the previous one-step backup to a two-step backup, updating the values of both *a_k_* and the action replayed previously; and recursively for the *n* + 1st step continuing an *n*-step backup conducted up til now. Note that *n*-step backups are only valid if the choices, after the first, are on-policy with respect to the target function *Q*\* (this is why only the maximal action constitutes a continuation).

Such sequence-extending experience activations permit a special learning step, and a corresponding special case of need/gain computation. If a sequence-extending experience is activated, the corresponding learning rule applies an *n*-step Bellman update at each of the preceding states in the sequences (i.e. it updates the value of all *n* preceding state/actions according to their subsequent cumulative, discounted rewards over the whole trajectory, plus the *Q*-value of the best action *a* at the added state *s′_k_*.) Implementationally, this can be accomplished using a *Q*(1) update rule over eligibility traces that are cleared whenever a sequence is *not* continued. The gain for this update, then, accumulates the gain over each of these state updates according to any policy changes at each, and this sum is multiplied by the need for the last state *s′_k_*. Altogether, this change allows the model to consider the set of one-step sample backups in addition to extending the previous *n*-step backup into a *n* +1-step backup, though their *EV B* is assessed myopically at each step.

The agent was allowed 20 planning steps at the beginning and at the end of each episode. Because the gain term is a function of the current set of *Q*-values, the utilities *EV B* were re-computed for all experiences after each planning step. In order to ensure that all 20 planning steps were used, a minimum gain of 10^−10^ was used for all experiences.

Prior to the first episode, the agent was initialized with a full set of experiences corresponding to executing every action in every state (equivalent to a full state-action-state transition model, which in sparse environments like these can be inferred directly from visual inspection when the agent first encounters the maze), including transitions from goal states to starting states. The state-state transition probability model 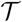 (for the need term) was initialized from this model under a random action selection policy, and thereafter updated after each transition using a delta rule with learning rate 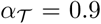. In all simulations in the online setting, the need term was then estimated from the SR matrix, 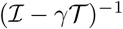. In the only simulation of sleep replay (Fig. 4g,h), where the agent is not located in the environment where need is computed, we estimated the need term as the stationary distribution of the MDP, i.e., the vector *µ* such that 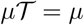.

### 4.3 Formal derivation

Below is a formal derivation of EVB for the general case of stochastic environments. Let the agent be in state *S_t_* = *s* at time *t*. The expected return from following policy *π* is defined as 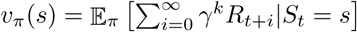, and the true (yet unknown) value of taking action *a* in state *s* and following policy *π* thereafter is denoted by *q_π_*(*s, a*). The utility of updating the agent’s policy from *π_old_* to *π_new_* is:

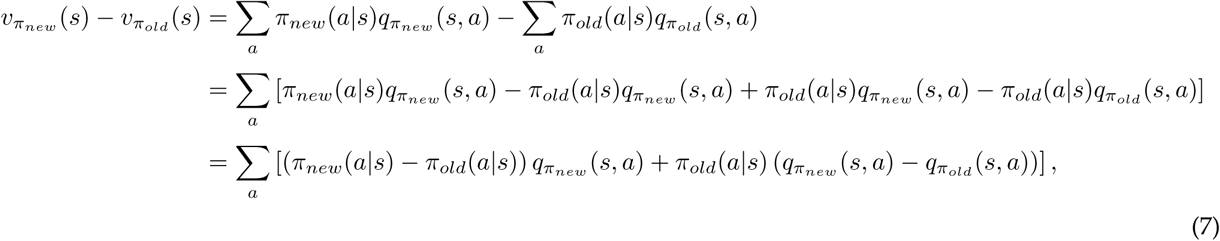
where we have both added and subtracted the term 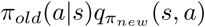 on the second line.

We then write *q*(*s, a*) in terms of *v*(*s*) using the definition of the MDP dynamics, 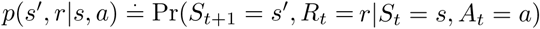:

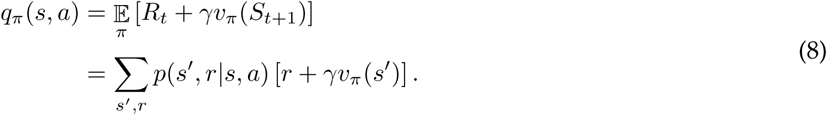

Since the MDP dynamics does not depend on *π*, we can write:

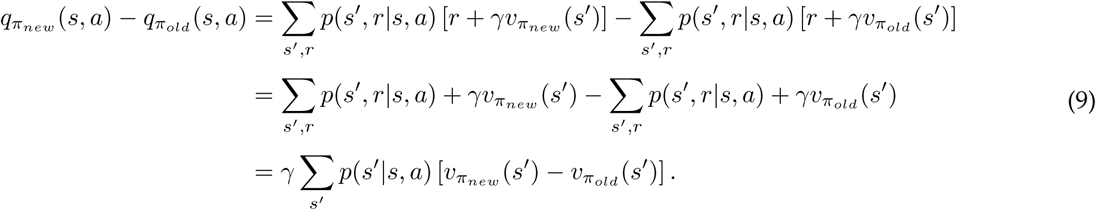

Substituting this result on (7):

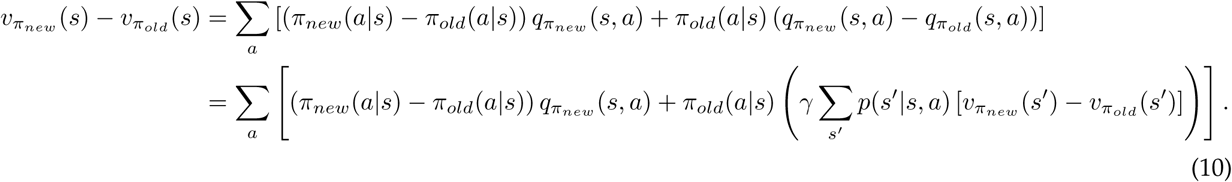

Notice that (10) contains an expression for 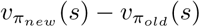 in terms of 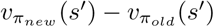. We can use this to ‘unroll’ the expression and write 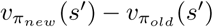 in terms of 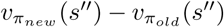. After repeated unrolling we obtain:

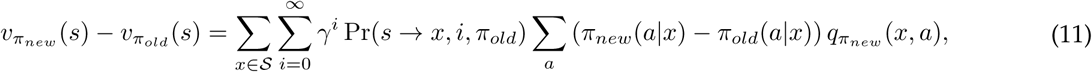
where Pr(*s* → *x*, *i*, *π_old_*) is the probability of transitioning from state *s* to state *x* in *i* steps under policy *π_old_*.

Since the effect of a backup on state-action pair (*s_k_,a_k_*) is localized at a single state for punctate representations, *π_new_*(*a*|*s_i_*)= *π_old_*(*a*|*s_j_*), ∀*i, j* ≠ *k*, and thus there is only one non zero term on the first summation:

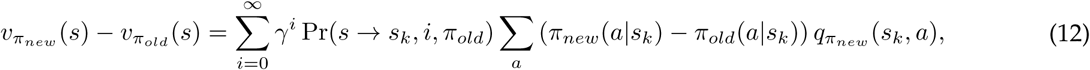

Denoting 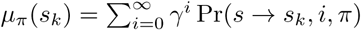 as the discounted number of time steps in which *S_t_* = *s_k_* in a randomly generated episode starting in *S_t_* = *s* and following *π*, we have:

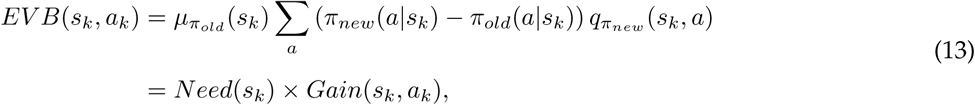
where 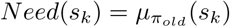 and 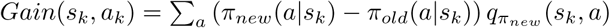.

We note that the same framework can be readily extended to the function approximation case and to policy gradient methods — i.e., computing the utility of a policy change even when the policies differ in multiple states (by using equation (11)). In this more general case, the above derivation corresponds to a discrete version of the Policy Gradient Theorem^74^.

### 4.4 Code availability

All simulations were conducted using custom code written in MATLAB v9.1.0 (R2016b). All results presented in this manuscript can be replicated by custom code available upon request.

## Acknowledgements

We thank Máté Lengyel, Daphna Shohamy, and Daniel Acosta-Kane for many helpful discussions, and Dylan Rich for his comments on an earlier draft of the manuscript. We acknowledge support from NIDA through grant R01DA038891, part of the CRCNS program, and Google DeepMind. The content is solely the responsibility of the authors and does not necessarily represent the official views of any of the funding agencies.

## Author contributions

Conceptualization, M.G.M. and N.D.D.; Methodology, M.G.M. and N.D.D.; Software, M.G.M.; Simulations, M.G.M.; Writing – Original Draft, M.G.M. and N.D.D.; Writing – Review & Editing, M.G.M. and N.D.D.; Funding Acquisition, N.D.D.

## Author information

The authors declare no potential conflict of interest.

